# Intermittent protein restriction elevates food intake and plasma ghrelin in male mice

**DOI:** 10.1101/2024.03.01.582931

**Authors:** K.L Volcko, H. Taghipourbibalan, J.E. McCutcheon

## Abstract

Low-protein diets affect body weight, body composition, food intake, and food preferences in mice. Furthermore, single periods of protein restriction can have lasting effects on these parameters. We sought to examine the effect of multiple, short, bouts of protein restriction, relative to long-term maintenance on either a control (NR) or protein-restricted (PR) diet. We found that male mice experiencing intermittent protein restriction (IPR) were indistinguishable from NR mice in terms of body weight and composition, but had food intake and plasma ghrelin as high as mice on PR diet, even when they were returned to control diet. This was not found in female mice. The results of this experiment highlight the importance of diet history on food intake and ghrelin levels in male mice, and the difference in how PR diet might affect male and female mice.

## 1. Introduction

Low-protein diets supplying 4-7% of caloric demands from protein have wide-ranging effects on physiology and ingestive behavior. In male mice, low-protein diets often lead to a reduction in body weight, relative to mice eating diets that are sufficient in protein (Fontana et al., 2016; Green et al., 2022; Hill et al., 2019, 2022; Laeger et al., 2014; Larson et al., 2017; Trautman et al., 2023; Wu et al., 2021) and, interestingly, the reduction in body weight occurs despite increased food intake (Fontana et al., 2016; Green et al., 2022; Hill et al., 2019; Laeger et al., 2014; Trautman et al., 2023; Volcko & McCutcheon, 2022; Wu et al., 2021). This mismatch appears due to increased energy expenditure observed in mice on low-protein diets (Hill et al., 2019, 2022; Wu et al., 2021). Low-protein diets also affect body composition. Some studies report a reduction in both fat and lean mass (Fontana et al., 2016; Hill et al., 2022; Larson et al., 2017; Trautman et al., 2023; Wu et al., 2021) and some in only lean mass (Green et al., 2022). Still others have found increases in fat mass (Simpson et al., 2013; Sørensen et al., 2008), though it is worth noting these studies have had 9% protein as their lowest protein levels, unlike the other studies cited which ranged from 4 – 7%; diets containing more than 10% protein seem to be above the threshold for inducing changes in male mice (Wu et al., 2021). Female mice have been studied far less frequently than male mice, but low-protein diets do not seem to have as large of an effect, if any, on body weight or body composition of female mice (Blais et al., 2018; Green et al., 2022; Larson et al., 2017; Volcko & McCutcheon, 2022); some studies do not find an effect of low-protein diet on food intake in female mice (Green et al., 2022; Larson et al., 2017) while others find increased intake (Blais et al., 2018; Volcko & McCutcheon, 2022).

Food choice is different in mice eating diets sufficient in protein compared to those eating a low-protein diet. Specifically, low-protein diets cause rodents to prefer sources of protein, such as the milk protein casein, over sources of carbohydrate, such as maltodextrin (Chiacchierini et al., 2021; Hill et al., 2019; Murphy et al., 2018; Volcko & McCutcheon, 2022). Despite differences between males and females in metabolic responses to low-protein diets, we have recently shown that this heightened protein preference is nearly identical in male and female mice (Volcko & McCutcheon, 2022).

The effects of low-protein diets can persist long after the animals are eating a normal diet again. For example, offspring of dams fed a low-protein diet during pregnancy and lactation have higher food intake even at a year of age (Qasem et al., 2016), and rats fed a low-protein diet during adolescence or early adulthood and a normal-protein diet thereafter, show elevated food intake and higher fat mass in adulthood than rats who did not experience a period of protein restriction (de Oliveira et al., 2013; Malta et al., 2014). The same group, however, did not find an effect when the diet manipulation occurred later in adulthood (Malta et al., 2016), indicating that the age of the rats was critical in determining the long-term effects of a period of protein restriction.

If a single period of protein restriction, in some developmental windows, can affect later behavior and body composition, one may also wonder if repeated bouts of protein restriction have an effect. This was explored in a recent study by Torres et al. (2022), in which individual male mice alternated between control and low-protein diets, with varying times on each diet (from 3 - 9 weeks). This study found that mice regulated body weight and food intake depending on which diet they were eating (i.e., eating more but gaining less weight when on the low-protein diet compared to the control diet) (Torres et al., 2022). Because this was a single-case experimental design, however, it did not examine whether repeated periods of protein restriction would have a long-term effect on body weight and composition compared to if no restriction was experienced. We were curious to know what would happen to food intake, body weight and composition, and plasma ghrelin levels if mice alternated between control and protein-restricted diets every seven days, and how this would compare to mice with continuous access to either NR or PR diet.

## 2. Methods

### 2.1 Animals

The experiment was conducted on two cohorts of mice, with male mice and female mice run at separate times. As such, the first cohort consisted of forty-two male C57BL/6NRj mice, and the second cohort of thirty-six female C57BL/6NRj mice (all mice 6 – 8 weeks old on arrival to the facility). All mice were purchased from Janvier (France) and housed in a temperature- and humidity-controlled room, maintained on a 12:12 light:dark cycle (lights-on at 07:00). Two mice shared each cage, separated by a perforated divider that allowed visual, olfactory, and auditory communication but prevented major physical contact, and allowed individual measurements of food intake. Water and food were available *ad libitum*. A third of the mice were fed a control diet (20% casein by weight and 18% of calories from protein; D11051801; Research Diets), a third of the mice an isocaloric protein-restricted diet (5% casein by weight, and 4% calories from protein; D15100602; Research Diets), and a third of the mice alternated between the two diets. The three groups, each with 12-14 mice, were therefore the non-restricted controls (NR), the chronically protein-restricted (PR), and the intermittently protein-restricted (IPR). All mice were fed the control diet for a week at the start of the experiment (Fig 1A). Both mice from each cage experienced the same diet condition. All animal care and experimentation followed the EU directive 2010/63/EU for animal experiments, and was approved by the National Animal Research Authority in Norway.

**Figure 1:**
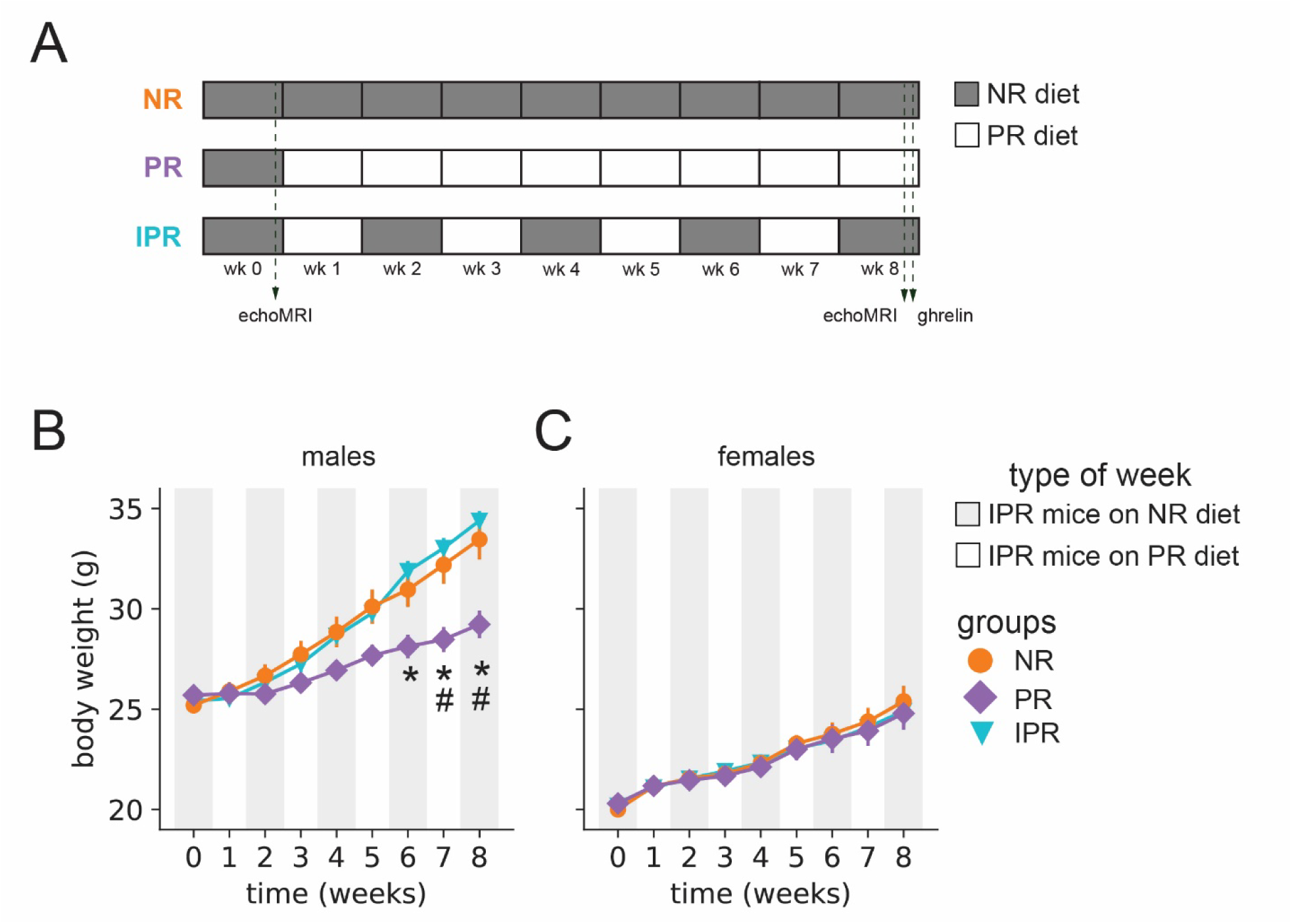
Schematic of experimental groups, and body weight. After a week of baseline measurements while on a normal protein diet (NR), some mice continued eating only this diet, some were given a protein-restricted diet (PR), and others alternated between the two diets (IPR) (A). Over the course of the experiment, body weight increased in male (B) and female (C) mice. Male mice on PR diet weighed less than IPR mice from week 6 and onwards, and less than those NR diet on week 8. Groups did not differ in body weight in female mice. Data are mean +/- SEM. Error bars too small to be visible. * PR differs from IPR, # PR differs from NR.

### 2.2 Body weight and food intake

Body weight was measured three days per week. For the male cohort, food intake was measured two days per week: one day immediately before the IPR group changed diets, and one day immediately after the IPR diets were switched. These two measurements were averaged for an average intake per week. For female cohort, food intake was measured weekly.

### 2.3 Body composition

Body composition was assessed at baseline (when all mice were on control diet) and after 8 weeks (at the end of the experiment). Mice were weighed early in the light phase, and body composition examined 0-6 hours later in an echoMRI-700 Body Composition Analyzer (echoMRI, Houston, TX, USA), with an equal number of mice from each diet condition measured earlier and later in the day.

### 2.4 Plasma levels of ghrelin

At the end of the experiment, plasma was collected, with half the mice in each diet group killed at each of two time points. In the “dark onset” condition, plasma was collected at the beginning of the dark phase / end of the light phase with food removed 4-6 hours beforehand. In the “light onset” condition, plasma was sampled at the beginning of the light phase / end of the dark phase. At the time of plasma collection, mice in the IPR group had switched back to NR diet 7 days earlier. Mice were killed under isoflurane anesthesia by cervical dislocation and decapitation before trunk blood was collected in K_2_ EDTA-coated tubes (16.444.100, Sarstedt) with 1 mg/ml AEBSF (Merck SBR00015) and placed on ice. Blood was centrifuged, within an hour of collection, for 10 min at 3000 rpm and 4°C (Mikro 220R centrifuge). Plasma was separated, acidified with hydrochloric acid to a concentration of 0.05N, and frozen at −70°C until analysis. Active ghrelin was assessed by ELISA (Millipore EZRGRA-90K) on a ClarioStar microplate reader (BMG Labtech).

### 2.5 Data analysis

Statistical analyses were conducted using JASP (https://jasp-stats.org/). All post-hoc tests were corrected for multiple comparisons using the Holm method. The study was not pre-registered. Data and code used to make figures are available at the following links: https://doi.org/10.5281/zenodo.13644179 and https://github.com/mccutcheonlab/intermittent-protein.

Body weight per week for each mouse was an average of three measurements. A mixed model ANOVA was conducted, with Group (NR, PR, or IPR) as between-groups factor and Time as within-groups factor. Body composition measures were mixed-design ANOVA with Group and Time (baseline vs end of experiment) as factors.

Food intake per week for each mouse was an average of two measurements for males, and a single weekly measurement for females. Values over 5 g per day, or under 1 g per day, were excluded as they were almost certainly erroneous. There were three occasions in which an estimated value for weekly food intake was used, so that we did not lose data for the entire mouse. In the female cohort, food intake for a single PR mouse for week 4 was estimated by averaging the weekly intake of this mouse on all other weeks (excluding baseline when she was eating NR diet). This was similar to the approach taken in the male cohort for a single NR mouse during week 2, though his baseline measurement was included since he had NR diet throughout. In the male cohort a PR mouse had its baseline intake estimated by using the average intake of all the other mice. The average food intake for each mouse on weeks when the IPR group was on NR, and when the IPR group was on PR, were compared between groups. A mixed model ANOVA was conducted, with Group as between-groups factor and Type of Week (IPR on NR diet, or IPR on PR diet) as within-groups factor. The average daily food intake per week was used to calculate average daily protein intake, and the same statistical analysis conducted as for food intake.

Plasma ghrelin was analyzed as a factorial ANOVA, with Group and Condition as factors. Due to the number of samples able to be run on a single ELISA plate, 3 female mice (1 NR, 1 PR, 1 IPR, all from the “light onset” condition) were not included. One mouse (male NR “dark onset”) was two standard deviations above the group mean and was therefore considered an outlier and removed from the analysis.

## 3. Results

### 3.1 Body weight

In males, body weight, as expected, increased over time as evidenced by a main effect of Time (Fig 1B; F_8, 312_ = 357.159, p < 0.001). There was also a main effect of Group (F_2, 39_ = 4.425, p = 0.019), as well as a Time x Group interaction (F_16, 312_ = 22.177, p < 0.001). Post-hoc tests on the interaction revealed that by week 6, IPR mice were significantly heavier than PR mice. Both IPR mice and NR mice groups were heavier than PR mice also at week 8. At no point did IPR mice and NR mice differ in body weight.

In females, body weight also increased over time (Fig. 1C; F_8, 264_ = 113.22, p < 0.001). There were no differences between the groups (F_2, 33_ = 0.047, p = 0.954), nor a Group x Time interaction.

### 3.2 Body composition

In males, fat mass increased between baseline and after eight weeks on experimental diets (Fig 2A; main effect of Time, F_1, 39_ = 305.706, p < 0.001). There was also a significant main effect of Group (F_2, 39_ = 22.91, p < 0.001), and a Group x Time interaction (F_2, 39_ = 9.813, p < 0.001). Post-hoc tests on this interaction indicated that although each of the groups had a higher fat mass at the end of the experiment than at the beginning, PR mice had significantly less fat mass at the end than either NR mice or IPR mice; NR and IPR mice did not differ from one another (Fig 2E). Analysis of lean mass also showed a significant main effect of Time (Fig 2C; F_1, 39_ = 50.109, p < 0.001), but no main effect of Group (F_2, 39_ = 2.178, p = 0.2). The Time x Group interaction was significant (F_2, 39_ = 8.5, p < 0.001), and post-hoc tests showed that only the NR mice and IPR mice gained lean mass over the eight weeks of the experiment (Fig 2G). At the end of the experiment IPR mice had a higher lean mass than did PR mice, but PR mice and NR mice did not differ in lean mass.

**Figure 2:**
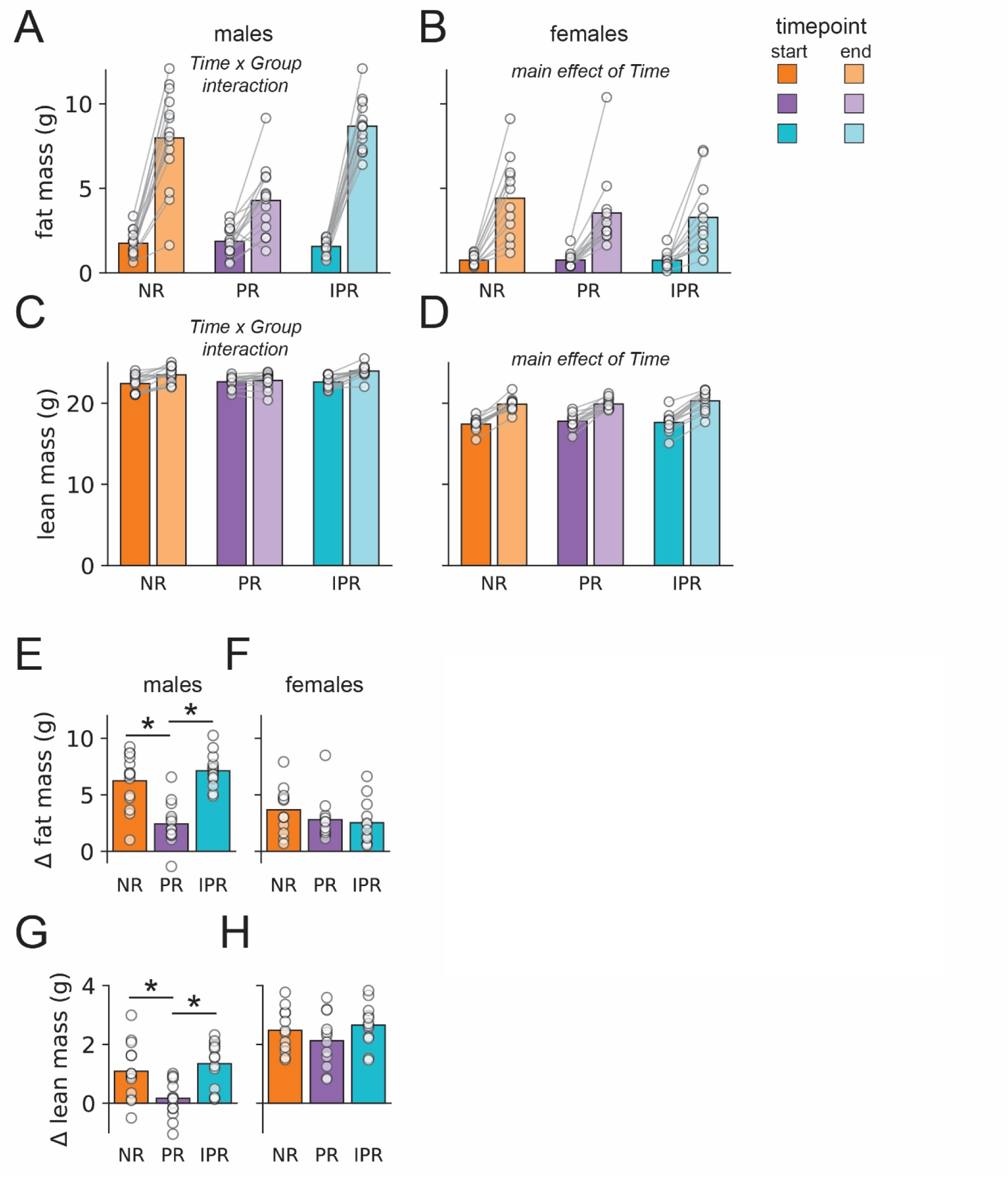
Body composition. Fat mass in male (A) and female (B) mice increased over time. Lean mass was only increased in NR and IPR males (C), but in all groups in female mice (D). Change in fat mass between start and end of the experiment was higher in NR and IPR males than PR males (E), with no differences between groups in females (F). Change in lean mass was also higher in NR and IPR males than PR males (G), with again no differences in females (H). Bars are mean and circles show individual data points. * p < 0.05 vs PR mice.

In females, fat mass increased over time (Fig 2B; F_1, 33_ = 79.768, p < 0.001) but there was no main effect of Group (F_2, 33_ = 0.644, p = 0.532) nor a Group x Time interaction (Fig 2F; _2, 33_ = 1.058, p = 0.359). A similar pattern of results was found for lean mass, such that lean mass was higher at the end of the experiment than at baseline (Fig 2D; F_1, 33_ = 329.289 p < 0.001), but there were no group differences (F_2, 33_ = 0.529, p = 0.722) nor Group x Time interaction (Fig 2H; F_2, 33_ = 1.359, p = 0.271).

### 3.3 Food intake

In males, average food intake consumed each week was different across time (Fig 3A; F_8, 312_ = 7.565, p < 0.001) and between groups (F_2, 39_ = 27.66, p < 0.001), and there was a significant Time x Group interaction (F_16, 312_ = 2.643, p < 0.001). Post-hoc testing revealed that NR males ate less food than did PR males on weeks 3, 4, 6, 7 and 8, and less than IPR mice did on weeks 3, 6, 7, and 8; IPR mice were eating NR diet on weeks 6 and 8. Furthermore, we compared the average intake of mice during weeks in which the IPR group was on NR diet or PR diets (Fig 3C). Here, we found that intake did not differ by type of week (IPR mice on NR diet vs. PR diet) (F_1, 39_ = 0.512, p = 0.512), but there was a significant main effect of Group (F_2, 39_ = 30.21, p < 0.001) and a Type of Week x Group interaction (F_2, 39_ = 3.433, p = 0.042). As expected, neither NR mice nor PR mice ate different amounts of their respective diets depending on whether IPR mice were eating NR or PR diet, but, interestingly, food intake also did not differ in IPR mice depending on which diet they were currently consuming. As such, regardless of which diet they were eating, IPR mice ate the same amount as PR mice and more than NR mice.

**Figure 3:**
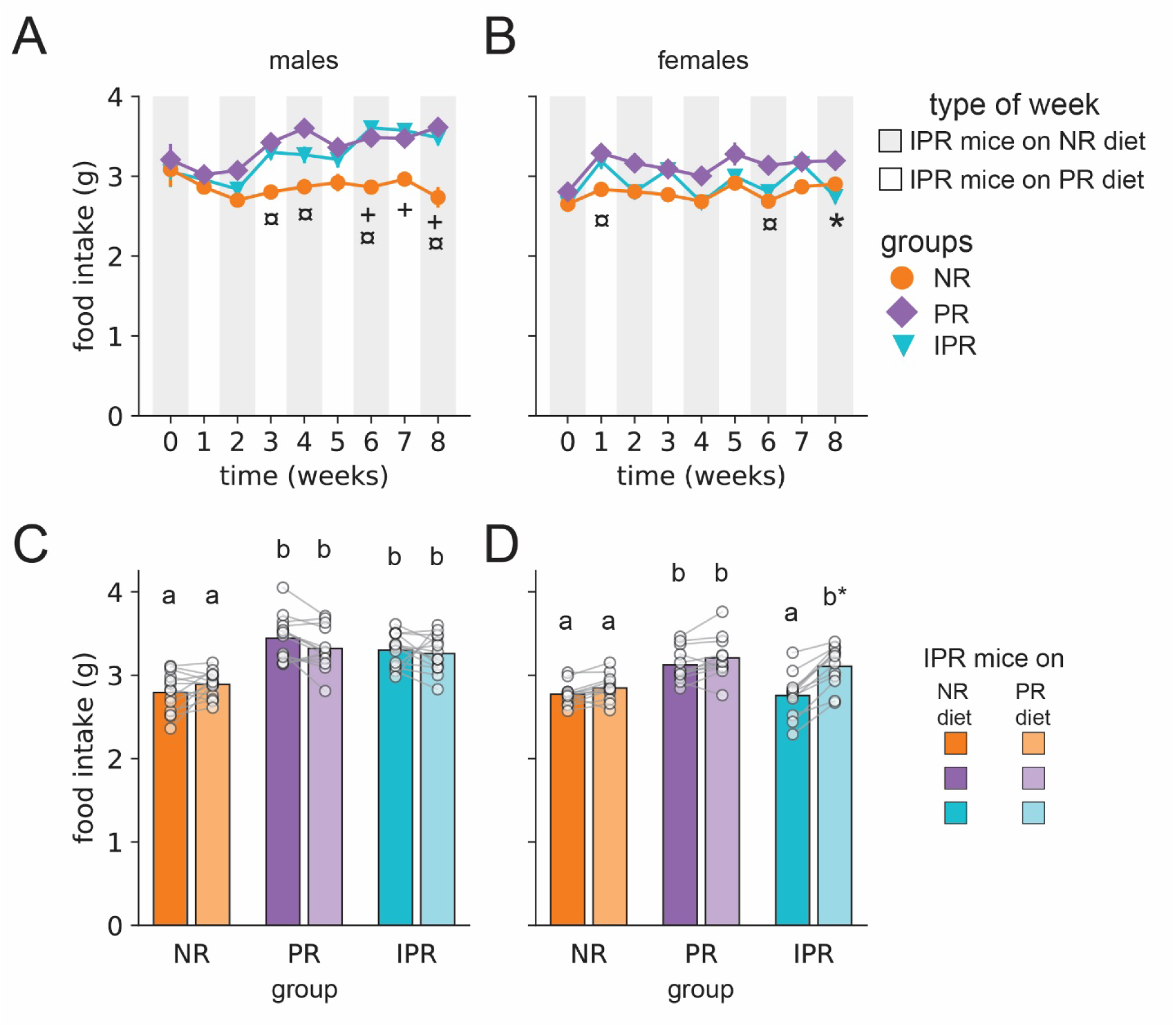
Food intake. Male PR mice ate significantly more than NR mice on weeks 3, 4, 6, 7 and 8; IPR male mice ate more than NR mice on week 3 and the last three weeks of the experiment (A). In female mice, PR mice ate more than NR on weeks 1 and 6, while IPR ate more than PR on week 8 (B). Average intake on weeks in which IPR was on NR diet or PR diet did not differ in any male groups, but PR and IPR males ate more than NR males regardless of week type (C). In female mice, IPR mice ate more when on PR diet than NR diet; their intake was well-matched to that of the mice on the chronic diets (D). In A and B, data are mean +/- SEM. Error bars are too small to be visible. In C and D, bars are mean and circles show individual data points. ¤ NR differs from PR, * PR differs from IPR, + IPR differs from NR, lower-case letters indicate p < 0.05 (e.g., a is different from b, b* p = 0.053 difference vs NR mice when IPR mice are on PR diet).

We also calculated average daily protein intake for each week. For male mice, there was a main effect of Time (F_8, 312_ = 137.077, p < 0.001) and Group (F_2, 39_ = 692.149, p < 0.001), and a Time x Group interaction (Fig 4A; F_16, 312_ = 91.986, p < 0.001). Post-hoc tests revealed no differences during week 0, the baseline week, when all mice were on NR diet, but thereafter NR males consumed significantly more protein than PR males did. When IPR males were on PR diet they ate as much protein as PR males, and when they were on NR diet they ate as much protein as NR males did, although in later weeks they tended to eat slightly more protein per day, a difference which only reached significance during week 6 and week 8. Average protein intake per day on weeks when IPR male mice were on NR diet or PR diet had main effects of Type of Week (F_1, 39_ = 1012.624, p < 0.001) and Group (F_2, 39_ = 1079.905, p < 0.001), and a Type of Week x Group interaction (Fig 4C; F_2, 39_ = 1098.825, p < 0.001). Post-hoc tests showed that PR mice consumed less protein than NR mice (as was expected due to the different protein content in their respective diets) and that there was no difference in protein intake by Type of Week for either NR mice or PR mice (also as expected since nothing changed between these types of weeks for the NR and PR groups). IPR male mice consumed a greater amount of protein than NR and PR mice, when they were on NR diet, and the same amount as PR mice did when they were on PR diet.

**Figure 4:**
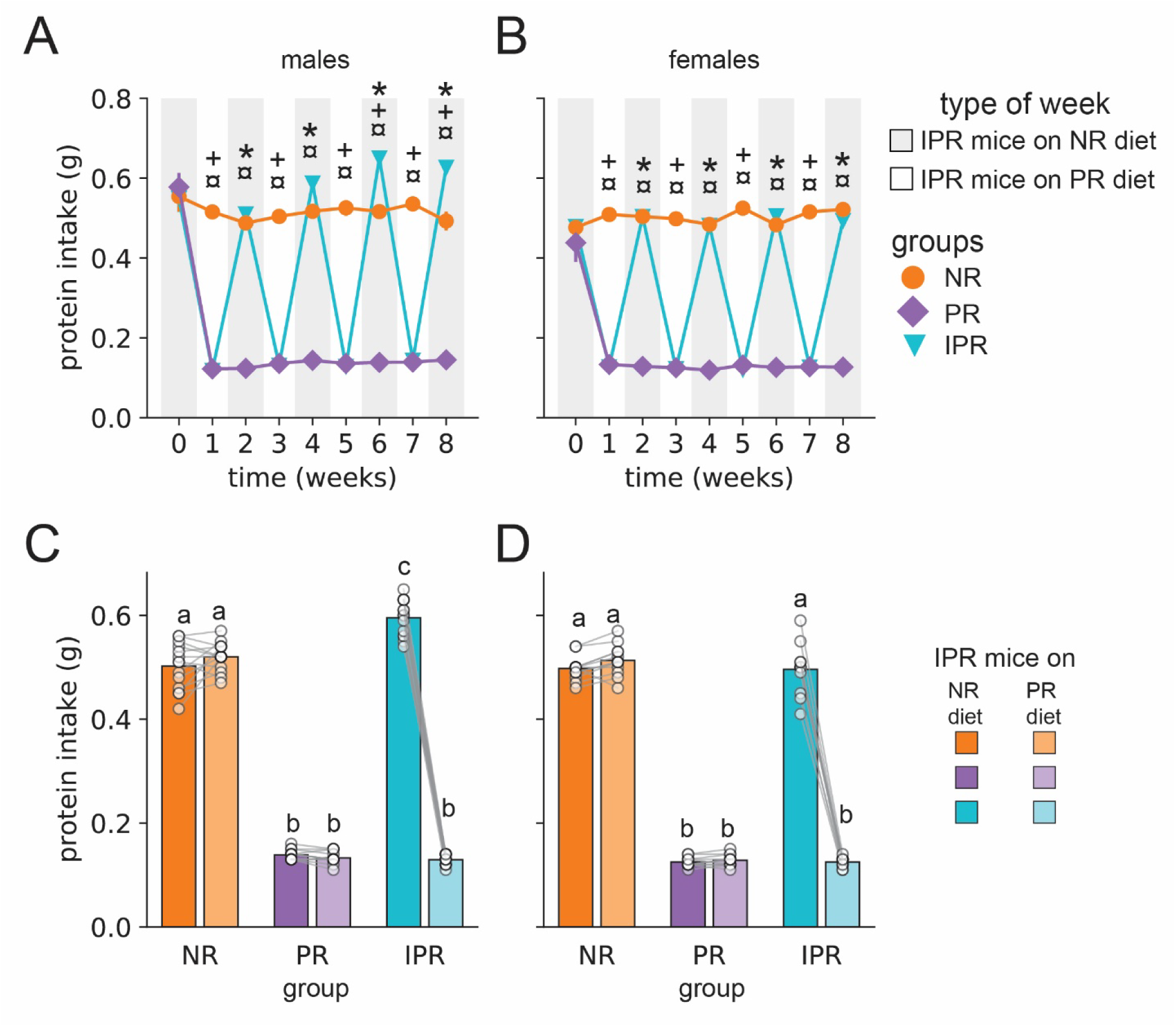
Protein Intake. Male NR mice ate more protein than male PR mice, with male IPR mice changing their protein intake depending on which diet they were on; on weeks 6 and 8 they exceeded the protein intake of NR mice (A). Female NR mice ate more protein than female PR mice, with IPR matching intake to the group whose diet they were eating (B). On weeks when IPR mice were on NR diet, male IPR mice ate, on average, more protein per day than the male mice on NR diet (C), while female IPR mice had equivalent protein intake as NR mice on weeks they were on NR diet (D). In A and B, data are mean +/- SEM. Error bars are too small to be visible. In C and D, bars are mean and circles show individual data points. ¤ NR differs from PR, * PR differs from IPR, + IPR differs from NR, lower-case letters indicate p < 0.05 (e.g., a is different from b).

In females, food intake by week had significant main effects of Time (Fig 3B; F_8, 264_ = 15.425, p < 0.001) and Group (F_2, 33_ = 8.503, p < 0.001), as well as a significant Time x Group interaction (F_16, 264_ = 3.05, p < 0.001). Post-hoc tests on the interaction revealed that NR mice and PR mice differed in intakes on weeks 1 and 6, while PR mice and IPR mice differed in intakes on week 8. Average intake on weeks when IPR mice were on PR diet or NR diet similarly had main effects of Type of Week (Fig 3D; F_1, 33_ = 66.354, p < 0.001) and Group (F_2, 33_ = 9.506, p < 0.001), and a Type of Week x Group interaction (F_2, 33_ = 18.108, p < 0.001). Post-hoc tests revealed that intakes of NR mice and PR mice remained stable (as expected), with NR mice eating less than PR mice. IPR mice, on the other hand, ate significantly more when on PR diet than on NR diet. When IPR mice were on NR diet their intake was the same as NR mice and lower than that of PR mice; when IPR mice were on PR diet their intake was the same as PR mice and there was a trend (p=0.053) towards their intake being higher than NR mice.

Average daily protein intake for females had a main effect of Time (F_8, 264_= 128.588, p < 0.001) and Group (F_2, 33_ = 581.555, p < 0.001), and a Time x Group interaction (Fig 4B; F_16, 264_ = 108.838, p < 0.001). Post-hoc tests revealed that all ate the same amount of protein at baseline. Following that, NR females and PR females differed in their average daily protein intake, and IPR females matched their intake to the group whose diet they were currently eating. Average protein intake per day on weeks when IPR mice were on NR diet or PR diet had main effects of Type of Week (F_1, 33_ = 821.072, p < 0.001) and Group (F_2, 33_ = 892.324, p < 0.001), and a Type of Week x Group interaction (Fig 4D; F_2, 33_ = 945.385, p < 0.001). Post-hoc tests showed that PR mice consumed less protein than NR control mice (as was expected due to their different diets) and that there was no difference in protein intake by Type of Week for either NR mice or PR mice. IPR female mice consumed an equivalent amount of protein as NR mice when they were on NR diet, and as PR mice when they were on PR diet.

### 3.4 Plasma ghrelin

In male mice, plasma ghrelin levels near dark onset and light onset were analyzed with repeated measures ANOVA (Fig 5). We found a main effect of Group (F_2, 35_ = 8.041, p = 0.001) and no effect of Condition (F_1, 35_ = 0.529, p = 0.472), nor an interaction (F_2, 35_ = 0.286, p = 0.753). Post-hoc tests revealed that both PR mice and IPR mice had higher plasma ghrelin than did NR mice.

**Figure 5:**
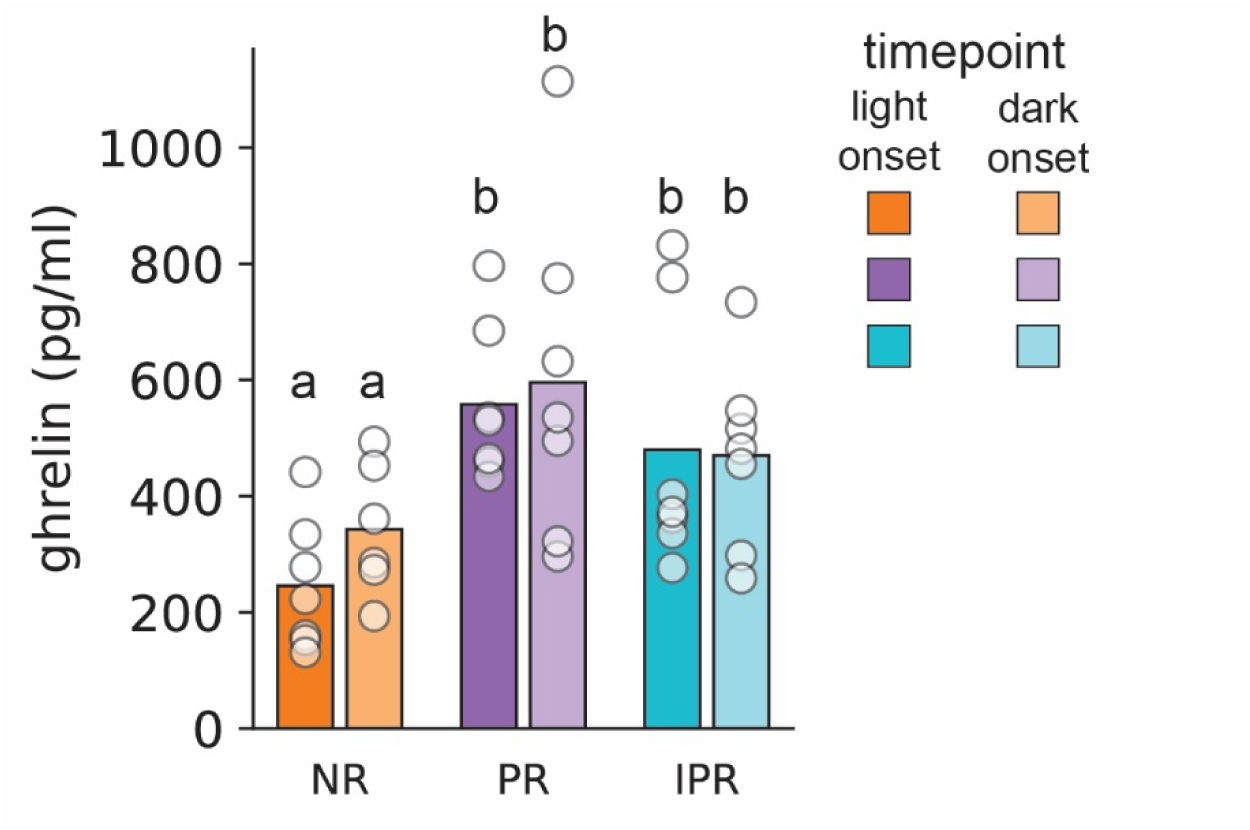
Plasma ghrelin in male mice. Males on PR diet, and IPR mice currently eating NR diet, both had higher plasma ghrelin than did NR mice. Bars are mean and circles show individual data points, lower-case letters indicate p < 0.05 (e.g., a is different from b).

In the female mice, many samples (9 out of 34) were out of range for the ELISA. Of these, all but one were from the light onset state. We therefore ran an ELISA only on the dark onset state, but found no effect of Group (data not shown; F_2, 14_ = 2.167, p = 0.151).

## 4. Discussion

Mice experiencing intermittent protein restriction did not show an intermediate phenotype between that of NR and PR mice. In male IPR mice, IPR mice were indistinguishable from NR control mice in terms of body weight and body composition. One reason could be that the average level of protein restriction was less in the IPR group than in the PR group. These mice spent half the time eating a 5% casein diet, and half the time eating a 20% casein diet, so on average were eating 12.5% casein. In terms of percent of calories from protein, the diets are 4% and 18%, respectively, which averages to 11% of calories from protein. This may be sufficient for maintaining growth, as it is above the 10% protein that may be a threshold for many parameters affected by protein restriction (Wu et al., 2021), although it is below the 13.6% that has been found to support growth and reproductive health in mice (Goettsch, 1960). This is important to consider because low-protein diets (and especially those low in certain amino acids) have been suggested as ways of improving metabolic health (for example, (Fontana et al., 2016; Rang et al., 2021). The present experiment highlights that at least some of these benefits may be contingent on long-term use of the diet or avoiding frequently switching between a low and normal protein diet. Female mice, regardless of group, did not differ in body weight or composition.

In male mice on low-protein diets, reduced body weight despite increased food intake is often found, and is explained by increased energy expenditure. These three consequences of low-protein diets are not always found, however. Some of this may be due to the length of the experiments. For example, previously in a shorter experiment we found increased food intake without changes in body weight (Volcko & McCutcheon, 2022). In the current experiment, body weight differences between NR and PR mice were only detected in the final, 8^th^ week of the diet manipulation. Another important consideration here is the housing method. Our mice were contact-housed, meaning they could communicate with a cage-mate but were physically separated and therefore unable to huddle together and share body heat. This has a large impact on their thermoregulation, and single-housed mice increase their energy expenditure and food intake relative to group-housed mice (Škop et al., 2021; Ziegler et al., 2022). Of particular interest for us is the possibility that the difference in energy expenditure between control and PR-fed mice was smaller than it would have been if the animals were group-housed, which in turn may have reduced diet-related differences. Several studies on protein restriction in mice have housed mice in pairs or groups (e.g. Green et al., 2022; Hill et al., 2022; Trautman et al., 2023), but many others have used single-housed mice as subjects (e.g. Hill et al., 2019; Laeger et al., 2014; Larson et al., 2017; Sørensen et al., 2008; Wu et al., 2021). Nonetheless, there could be a wide variation between facilities in terms of quantity and type of bedding provided, ambient temperature of the colony room, etc. All of these factors will impact thermoregulation and thereby energy expenditure, and a single-housed mouse may be more vulnerable to such perturbations than a group-housed mouse. In our experience, the male C57B6 mice are prone to fighting which has eliminated the option of group-housing, but it is an important consideration when viewing the data.

Interestingly, one measure in which intermittent protein restriction differed from control diet was in food intake. Protein restriction leads to higher caloric intake per day; this has been demonstrated many times in male mice (Fontana et al., 2016; Green et al., 2022; Hill et al., 2019; Laeger et al., 2014; Trautman et al., 2023; Volcko & McCutcheon, 2022; Wu et al., 2021) and is sometimes observed in female mice (Blais et al., 2018; Volcko & McCutcheon, 2022). In males, we found that the increase in food intake persisted even when the IPR male mice were consuming the control diet. Presumably, this persistent hyperphagia allowed the mice to compensate for a period when a reduced amount of protein was available. This is supported by the fact that on average, IPR males consumed more protein per day when on control diet than did the NR control males. Understanding precise parameters underlying this compensatory behavior – e.g., how persistent is the hyperphagia and, if a choice is available, is it directed towards certain foods? – will be important for understanding any consequences for human health and eating behavior. As such, future experiments should address these questions as well as consider which parts of the brain are altered by protein restriction to promote hyperphagia of both low- and normal-protein food sources. Importantly, a recent paper has reviewed the persistence of both behavioral responses and physiological parameters such as a change in neural activity (Soto & Morrison, 2024). The authors argue that behavior and physiology are dictated not solely by current nutritional requirements but also by historical nutritional conditions. Our results now support this notion, especially given that they were, on average consuming 12.5% casein (11% of calories from protein) and yet experiencing persistent heightened food intake, unlike mice maintained on 10% calories from protein which do not overeat (Wu et al., 2021).

Female mice on intermittent protein restriction did not show persistent hyperphagia, but were able to adapt their intake to whichever diet they were eating at the time. As others have pointed out, sex is an important factor in the effects of protein restriction (Green et al., 2022). It is worth noting that food intake was measured differently in male and female mice: in males, two 24-hour measurement periods (one immediately before and one immediately after IPR changed diets) were averaged to form a mean daily intake per week, whereas in female mice total food consumed in a week was divided by 7 for a average daily intake. In female mice we did measure 24-hour intake on a few occasions, and correlated the intakes using both methods. The R^2^ value (0.72 and 0.78, on two separate weeks, p < 0.001 on both occasions) demonstrates that using two methods may add some variability, but the relationship is strong enough that we feel confident in the major interpretation of the data: male mice on IPR show persistent high levels of intake regardless of their diet, whereas female mice on IPR do not.

Ghrelin levels, surprisingly, did not differ by sampling time point – i.e., “dark onset” vs. “light onset” – in our experiment. In the female cohort many samples from the “light onset” state were below the detection threshold, indicating that condition (near dark vs. light onset) did affect ghrelin levels. In the male cohort this was not the case; although this could be a sex difference, it is also likely that the minor difference in timing was important. In females, blood was sampled at the end of the dark phase and at the end of the light phase; in males, blood was sampled at the beginning of the light phase and at the beginning of the dark phase. This is a small difference in clock time but perhaps a very significant one in ghrelin secretion. Ghrelin rises during fasting and falls quickly after re-feeding (Tschop et al., 2000), and rodents typically eat in the dark with food intake concentrated at the beginning and end of the dark phase (Strubbe & Van Dijk, 2002). In light of this, we chose the time points with the assumption that ghrelin would be high after a short experimentally-imposed fast and at the beginning of the dark phase when eating normally occurs, and would be lower at the beginning of the light phase after the mice had experienced *ad libitum* overnight access to their respective diets. We did not find this. Plasma ghrelin follows a circadian pattern of secretion with higher levels in the light than dark, and the peak 5 hours after lights-on (Bodosi et al., 2004), though a different study found the peak to occur immediately at lights-on (Bertani et al., 2010). In contrast to this, another experiment found peak ghrelin immediately at lights-off (Sánchez et al., 2004), with a further study that examined ghrelin for a few hours near dark onset finding the peak 30 min before lights-out (Drazen et al., 2006). In short, different experiments find slight differences in when ghrelin is at its maximum and minimum, indicating that it is sensitive to many factors.

Despite failing to see a time-of-day difference in ghrelin in our experiment, we did find a significant Group difference, and this is intriguing. In males, there was a main effect of Group with PR mice having higher plasma ghrelin than NR mice, similar to what is seen in rats (Chaumontet et al., 2018). IPR mice, interestingly, had as high plasma ghrelin as PR mice, despite blood sampling occurring a week after they had returned to NR diet. Whether or not this potential difference in ghrelin might underlie the persistence of increased food consumption seen in PR and IPR mice warrants further study. Future experiments should also assay other feeding-related hormones. Of particular interest would be fibroblast growth factor 21 (FGF21), which is elevated during protein restriction and required for many of the changes in physiology and behavior resulting from low dietary protein (Hill et al., 2019, 2022; Laeger et al., 2014, 2016). The IPR mice in our experiment were, on average, consuming an adequate amount of protein, but alternating between low and normal levels in the diet. As little as 4 day of protein restriction dramatically elevates plasma FGF21 (Laeger et al., 2014), so the IPR mice in our experiment presumably had elevated FGF21 for at least several days in a row. How quickly FGF21 levels fall after returning to control diet, and whether or not this might contribute to the persistent hyperphagia would be interesting to know. Furthermore, because food intake in IPR males remained high regardless of which diet they were currently eating, it could be useful to examine if these periods of protein restriction caused epigenetic modifications that affected appetite.

Female mice on protein-restricted diet were remarkably similar to their controls eating a diet with sufficient protein, in all measures but food intake. This is striking when viewed in light of their protein-seeking behavior. Although we did not measure protein preference in this experiment, we have previously shown that female mice have as robust a shift in macronutrient preference as do male mice experiencing protein restriction (Volcko & McCutcheon, 2022). The lack of a change in body weight or composition in female mice on this diet indicates that they are able to obtain enough protein for growth by increasing their intake of protein-restricted diet, while males are not able to do so. One might speculate that this would lead to a greater drive to consume protein in males than females; in a sense, a male mouse can perhaps be seen as experiencing a higher degree of restriction on the same diet, because its protein requirements are higher. And yet, we do not see a commensurate increase in protein appetite. This is intriguing and suggests that the relationship between motivation to consume protein, and protein need, is not entirely linear.

Although we believe that the present experiments have yielded several interesting results, it is important to highlight the limitations of our study. A major limitation is that male and female mice were not tested in the same cohort, and there were some differences in how the experiments were conducted (e.g., how food intake was measured). This makes direct comparisons between the sexes impossible. The ghrelin measures did not show a time-of-day difference and the female plasma did not produce usable data, and caution is therefore advised when considering these findings. Moreover, many questions remain unanswered, such as whether IPR males would eventually exceed NR males in body weight, and how long their hyperphagia would last if kept on NR diet for longer than a week.

In summary, here we add support to earlier studies showing that male and female mice appeared to respond differently when challenged with a low-protein diet with respect to body weight, body composition, and food intake. Moreover, male mice completely compensate for repeated periods of low-protein diet exposure by increasing food intake and thereby protein intake, both while on the PR diet and for at least a week after return to a diet with a normal protein level.

A higher level of plasma ghrelin may be at least partially involved in this hyperphagia, although a causal link was not studied here.

## Acknowledgements

The authors wish to thank the excellent animal care staff at UiT; Neoma Boardman for use of a centrifuge; and Mette Kongstorp for helping with an ELISA.

## Author contributions

**Volcko, K.L**: Conceptualization, Investigation, Writing – Original Draft. **Taghipourbibalan, H**: Investigation. **McCutcheon, J.E**: Conceptualization, Supervision, Writing – Review and Editing. All authors have approved this manuscript.

## Declaration of Interest

None

## Funding

This work was supported by a Tromsø Research Foundation Starting Grant to JEM (19-SG-JMcC).

## Notes

https://doi.org/10.5281/zenodo.13644179

https://github.com/mccutcheonlab/intermittent-protein

### Competing Interest Statement

The authors have declared no competing interest.

### Summary of Updates

Revised Introduction and Discussion section. Figure 1 altered to include different axes. Extra figure on protein intake added (Fig. 4)

